# Neuroprotective Effect of Intraperitoneal Humanin-G in Retinal Degeneration of Royal College of Surgeons Rats

**DOI:** 10.64898/2026.03.20.713049

**Authors:** Bin Lin, Kevin Schneider, Mustafa Ozgul, Narcisa Ianopol, Magdalene Seiler

**Author notes:** Equal contribution (Bin Lin, Kevin Schneider). **Corresponding author:** Magdalene J. Seiler, Ph.D. Professor in Residence, Dept. Physical Medicine & Rehabilitation, Sue & Bill Gross Stem Cell Research Center, University of California, Irvine, 845 Health Sciences Road, 2035 Gross Hall Irvine CA 92697-1705.

## Abstract

This study aimed to examine whether Humanin-G (HNG), a mitochondrial derived peptide with cytoprotective properties, could improve the retinal function and gene expression profiles after intraperitoneal injections to Royal College of Surgeons (RCS) rats with Retinal Pigment Epithelium (RPE) dysfunction and retinal degeneration.

Starting at postnatal day 21 (p21), RCS rats received twice a week intraperitoneal injections of either Low Dose HNG (0.4 mg/kg), High Dose HNG (4mg/kg), or sham-saline for 1 or 4 weeks. Visual function was tested with full field scotopic & photopic electroretinography (ERG) and optokinetic testing (OKT) 1 and 4 weeks after first injection (WAFI). The rats were euthanized after the ERG and OKT (1 or 4 WAFI) and the dissected retinas and RPE were collected for RNA, cDNA and Quantitative Real-time PCR (qRT-PCR) analysis.

The results of our study showed that high dose (4mg/kg) HNG at 4 WAFI was associated with the largest change in gene expression in the RPE and retina of treated animals, altering expression of genes involved in apoptosis, oxidative stress, inflammation and retinal/RPE function. Analysis of a and b waves from scotopic and photopic ERG showed no difference between either low or high dose of HNG and sham injection at 4 WAFI. However, at 4 WAFI, the visual acuity in rats treated with high dose HNG showed significant improvement as compared to the rats treated with low dose of HNG or saline.

Most significantly, our findings support that HNG administered IP can modulate RPE/neuroretina cells and improve vision, thus may be a potential treatment for retinal degeneration diseases.

## Introduction

Mitochondrial DNA (mtDNA) (inherited maternally) can be classified into haplogroups, according to the accumulation of single nucleotide polymorphisms (SNPs). Mitochondria, with their small, but significant mtDNA, play important roles in human diseases, even when the disease is not caused directly by mitochondrial mutations ^1,2^. mtDNA variants can exert major influences on cellular energy pathways as well as mediate expression of nuclear genes related to complement, inflammation, apoptosis, autophagy, methylation and cell signaling transduction pathways ^3–11^.

Mitochondrial Derived Peptides (MDPs) are biologically active, short peptides (20-27 AAs), which are encoded from short open reading frames (sORF) in the human mtDNA ^12–15^. Humanin (HN), a 24-amino acid MDP, has been shown in vitro and in vivo to have anti-apoptotic, neuro-protective properties supporting cell survival ^14–17^. Circulating HN levels decrease with age and are associated with age-related diseases in mice, rats and humans ^18,19^. Furthermore, offspring of centenarians have higher HN levels compared to the aging population^18^. Altogether, our findings suggest that retaining HN levels with age may promote healthy aging. HN has been shown to be cyto-protective in Alzheimer’s disease, atherosclerosis, myocardial and cerebral ischemia, and Type 2 diabetes ^17,20^. HN and SHLP2 (Small Humanin-Like Peptide 2) can lower plasma amino acids and lipid metabolites associated with metabolic aging disorders ^21^.

In vitro studies have shown protective effects of HN against hypoxia-induced toxicity in RGC-5 cells ^22^. Pre- or co-treatment with HN protected SH-SY5Y neuroblastoma cells from toxicity induced by silver nanoparticles (AgNPs) ^23^. The HNG peptide, the HN analogue, improved mitochondrial function and decreased cell death in PC12 cells stressed with amyloidβ25-35 peptides ^24^. In human ARPE-19 retinal cybrids, HNG reduced amyloid-β induced cell stress ^25^. HNG has a glycine replacing serine at position 14, making it 1000-fold more potent in its protective functions.

An in vitro animal study showed rescue of cortical neuron viability after NMDA damage following 10 µmol/L HN treatment ^26^. In vivo, HN-treated diabetic mice demonstrated improved glucose tolerance and lower pancreatic-beta cell apoptosis due to activation of the Stat3 pathway ^27^. HNG treatment improved the cognitive functions in aging C57B1/6N mice and reduced the levels of inflammatory markers ^28^. While studies have shown HN or HNG improve retinal cells in vitro ^16,29,30^, there are no studies showing their in vivo effects in retinal degeneration models.

The Royal College of Surgeons (RCS) rat is an established model for retinal degeneration. This model has dysfunctional RPE due to a MerTK mutation leading to the accumulation of outer segment debris and photoreceptor death^31^, is associated with ER stress ^32^, and activation of mitochondrial and cytosolic calpain ^33^. The RCS model has been frequently used in the preclinical testing of drugs and stem cells for retinal degeneration ^34–40^. Recent studies have identified early mitochondrial dysfunction in RCS rats ^41^ and mtDNA deletions in the late stages of RCS retinal degeneration (>200d)^42^.

The present study investigates the effects of Humanin-G (HNG) intraperitoneal (IP) injections on the Royal College of Surgeons rat. We used intraperitoneal injections to deliver either low dose (0.4 mg/kg) or high dose (4mg/kg) HNG to the RCS rats and assessed the molecular changes in RPE cells and neuroretina, as well as vision change at 1 WAFI and 4 WAFI.

## Materials and Methods

### HNG Peptide Preparation

The custom synthesized HNG peptide with guaranteed TFA removal services was purchased from Genscript Inc. (Piscataway, NJ) and diluted with sterile 0.9% saline before the injection. PLGA (Poly(lactic-co-glycolic acid)) and PVA (Polyvinyl alcohol) polymers were purchased from Akina Inc (West Lafayette, IN). Methylene Chloride was purchased from Sigma-Aldrich (St. Louis, MO). Acetonitrile, HPLC water, LC-MS water, and formic acid were purchased from Fisher Scientific (Waltham, MA). Analytical grade solvents were used in all experiments.

### HNG Treatments for RCS Rats with Retinal Degeneration

#### Experimental animals

For all experimental procedures, animals were treated in accordance with the NIH guidelines for the care and use of laboratory animals, the ARVO Statement for the Use of Animals in Ophthalmic and Vision Research, and under a protocol approved by the Institutional Animal Care and Use Committee (AUP-17-097 and AUP 20-055). Non-nude rats of the immunodeficient Royal College of Surgeons (RCS) rat strain with the MERTK mutation (dysfunctional RPE) were used for the experiments ^39^.

#### Intraperitoneal Injection of HNG

Starting at postnatal day 21 (p21), rats were given an intraperitoneal injection of either “Low Dose” (0.4mg/kg) HNG; “High Dose” (4mg/kg) HNG; or sham-saline. Injections using either 22g or 27g needles were repeated twice weekly. The experiments were concluded after either 1 WAFI (short-term study) or 4 WAFI (long-term study) after the ERG and OKT tests were completed.

#### Full field Scotopic & Photopic Electroretinography (ERG)

For low dose injection, the rats were tested with ERG, using a Rodent ERG system (Diagnosys Celeris), at 4 WAFI. For high dose injection, the rats were evaluated for changes in visual function at 1 WAFI and 4 WAFI. After overnight dark-adaptation, rats were anesthetized by Ketamine/Xylazine (40-55 mg/kg Ket, 6 – 7.5 mg/kg Xyl), then the eyes were treated with 0.5% tetracaine (Bausch & Lomb) and 1% atropine eye drops (Akorn Pharmaceuticals, Lake Forest, IL). and then anesthesia was maintained by 1% isoflurane. The rats were placed onto a heating pad on a rodent exam table and positioned in front of a monocular mini-ganzfeld photostimulator. After applying GenTeal Lubricant Eye Gel, electrodes were placed onto the corneal surface. Visual responses with scotopic and photopic stimuli were recorded for both eyes, to obtain simultaneous recordings of the same animal using established protocols.

#### Optokinetic Testing (OKT)

For low dose injection, the rats were tested with OKT at 4 WAFI. For high dose injection, 1 and 4 weeks after the first injection of HNG (low dose and high dose) or saline, the visual acuity of RCS nude rats was measured by recording videos of optomotor responses to a virtual cylinder with alternating black and white vertical stripes (Optomotry, Cerebral Mechanics Inc., Alberta, Canada). The testing was described previously^36,43^.

Rats were dark-adapted for at least 1 hour prior to testing. Optomotor responses were recorded at 6 different spatial frequencies for one minute per frequency by testers blinded to the experimental condition. Both the left and right eyes were tested by alternating the direction of the moving stripes. Two independent tests were performed at each time point, with at least one hour time in between; one test going from lowest to highest frequency, and the other from highest to lowest frequency. The best visual acuity of the two tests was used for analysis. All tests were video recorded and evaluated off-line by two independent observers blinded to the experimental conditions. Any discrepancies between the two observers resulted in re-analysis of videos by a third observer, and data discussion before giving a final score, and prior to decoding the experimental condition.

#### Isolation of tissue, RNA and Amplification of cDNA

Upon removal of the anterior segment of the eye including the cornea, iris and lens, the retina was removed and immediately snap frozen in liquid nitrogen. The RPE was isolated by utilizing the “Simultaneous RPE cell Isolation and RNA Stabilization” (SRIRS method) ^44^. The posterior eye cup including the choroid, sclera and RPE was removed and immediately placed in a microcentrifuge tube containing 400µl of RNAprotect cell reagent (Qiagen). After a minimum of 10 minutes, the tube was agitated and RPE cells observed to be released into solution. The eye cup was then removed and the dissociated RPE cells pelleted. This provided selective isolation of the RPE and protection against RNA degradation in one step. RNA was isolated from RPE cells and the neuroretina using the PureLink RNA mini kit. cDNA was produced using SuperScript IV VILO (ThermoFisher) according to the manufacturer’s instructions.

#### Quantitative Real-time PCR (qRT-PCR) Analyses

qRT-PCR was performed on individual samples using PowerUp SYBR Green Master Mix (ThermoFisher Scientific) on an Applied Biosystems QuantStudio 5 Real-Time PCR system real time quantitative PCR detection system. Primers (QuantiTect Primer Assay (Qiagen) or KiCqStart Primers (Sigma)) were used to analyze 24 different genes: Inflammation and oxidative stress (*Ddit3, HSP*α*5, Il6, Il1*β*, Tnf*α*, Ccl2*); Apoptosis (*Bax, Casp3, Casp7, Casp9, Bcl2l1, Bcl2l13*); Antioxidation (*Sod2*); Photoreceptor markers (*Crx, Gngt1, Nrl, Rom1, E2f1*); and RPE markers (*Best1, Rlbp1, Rpe65, Tjp1*) (**Table 1**). Primers were standardized with the HMBS housekeeping gene. All analyses were performed in triplicate. The fold values were calculated using the 2^(-ΔΔCt) formula. All fold value changes of the HNG-IP treatments are calculated compared with saline control.

**Table 1.** Description of Genes Analyzed by qRT-PCR.

| Symbol | Gene Name | Gene RefSeq Number | Functions |
| --- | --- | --- | --- |
|  | **<u>INFLAMMATION PATHWAY</u>** |  |  |
| ***Ddit3*** | DNA-damage inducible transcript 3 | NM_001109986 | This gene encodes a member of the CCAAT/enhancer-binding protein (C/EBP) family of transcription factors. The protein functions as a dominant-negative inhibitor by forming heterodimers with other C/EBP members and preventing their DNA binding activity. The protein is implicated in adipogenesis and erythropoiesis, is activated by endoplasmic reticulum stress, and promotes apoptosis. Fusion of this gene and FUS on chromosome 16 or EWSR1 on chromosome 22 induced by translocation generates chimeric proteins in myxoid liposarcomas or Ewing sarcoma. Multiple alternatively spliced transcript variants encoding two isoforms with different length have been identified. |
| ***Il6*** | interleukin 6 | NM_012589 | Enables cytokine activity and interleukin-6 receptor binding activity. Involved cellular response to lipid; positive regulation of cell communication; and response to peptide hormone. Located in cytoplasm and extracellular space. |
| ***Il1b*** | interleukin 1 beta | NM_031512 | Enables cytokine activity. Involved in positive regulation of intracellular signal transduction; regulation of gene expression; and regulation of neurogenesis. Located in extracellular space. |
| ***Tnf*** | tumor necrosis factor | NM_012675 | Enables tumor necrosis factor receptor binding activity. Involved in positive regulation of cell communication; positive regulation of macromolecule metabolic process; and positive regulation of neuron death. Located in several cellular components, including external side of plasma membrane; extracellular space; and neuronal cell body. |
| ***Hspa5*** | heat shock protein family A (Hsp70) member 5 | NM_013083 | Enables misfolded protein binding activity and unfolded protein binding activity. Involved in cellular response to cAMP; cellular response to gamma radiation; and cellular response to metal ion. Acts upstream of or within response to endoplasmic reticulum stress. Located in membrane; mitochondrion; and smooth endoplasmic reticulum. |
| ***Ccl2*** | C-C motif chemokine ligand 2 | NM_031530 | Enables chemokine activity and heparin binding activity. Involved in cellular response to cytokine stimulus; cellular response to lipid; and leukocyte migration. Located in several cellular components, including perikaryon; perinuclear region of cytoplasm; and rough endoplasmic reticulum. Colocalizes with C-fiber. |
|  | **<u>ANTIOXIDANT PATHWAY</u>** |  |  |
| ***Sod2*** | superoxide dismutase 2 | NM_017051 | Enables several functions, including identical protein binding activity; manganese ion binding activity; and superoxide dismutase activity. Involved in hydrogen peroxide biosynthetic process; negative regulation of membrane hyperpolarization; and positive regulation of hydrogen peroxide biosynthetic process. Located in mitochondrial nucleoid. |
|  | **<u>APOPTOSIS PATHWAY</u>** |  |  |
| ***Bax*** | bcl2 associated x | NM_017059 | Enables BH domain binding activity; chaperone binding activity; and heat shock protein binding activity. Involved in glial cell apoptotic process; mitochondrion organization; and response to corticosterone. Located in cytosol; mitochondrial outer membrane; and perinuclear region of cytoplasm. |
| ***Bcl2l1*** | bcl2-like 1 | NM_001033672 | Enables several functions, including GTPase binding activity; MDM2/MDM4 family protein binding activity; and cysteine-type endopeptidase inhibitor activity involved in apoptotic process. Involved in cellular response to cytokine stimulus; cellular response to lipid; and positive regulation of transport. Located in cytosol; mitochondrial outer membrane; and presynapse. Colocalizes with clathrin-coated pit; mitochondrial membrane; and synaptic vesicle membrane. |
| ***Bcl2l13*** | bcl2-like 13 | NM_001398849 | Over-expression leads to apoptosis. Mitochondria specific protein. |
| ***Casp3*** | caspase 3 | NM_012922 | Enables death receptor binding activity; phospholipase A2 activator activity; and protease binding activity. Involved in luteolysis; nervous system development; and positive regulation of amyloid-beta formation. Located in cytosol; membrane raft; and neuronal cell body. Part of death-inducing signaling complex. |
| ***Casp7*** | caspase 7 | NM_022260 | Enables cysteine-type peptidase activity. Involved in aging; leukocyte apoptotic process; and striated muscle cell differentiation. Located in cytosol and intracellular membrane-bounded organelle. |
| ***Casp9*** | caspase 9 | NM_031632 | Enables cysteine-type peptidase activity. Involved in glial cell apoptotic process; response to estradiol; and response to ischemia. Located in cytosol; mitochondrion; and nucleus. |
|  | **<u>PHOTORECEPTOR MARKERS</u>** |  |  |
| ***Crx*** | cone-rod homeobox | NM_021855 | Enables DNA binding activity. Involved in circadian rhythm and positive regulation of photoreceptor cell differentiation. Located in nucleus. |
| ***E2f1*** | E2F transcription factor 1 | NM_001100778 | Enables protein kinase binding activity and sequence-specific DNA binding activity. Involved in cellular response to hypoxia; cellular response to nerve growth factor stimulus; and positive regulation of glial cell proliferation. Located in cytoplasm and nucleus. |
| ***Rom1*** | retinal outer segment membrane protein 1 | NM_001009690 | Predicted to enable protein homodimerization activity; be involved in cell adhesion; act upstream of or within detection of light stimulus involved in visual perception; protein complex oligomerization; and retina development in camera-type eye. Predicted to be located in photoreceptor outer segment membrane and an integral component of plasma membrane. |
| ***Nrl*** | neural retina leucine zipper | NM_001106036 | Predicted to enable DNA-binding transcription activator activity, RNA polymerase II-specific; leucine zipper domain binding activity; and promoter-specific chromatin binding activity. Predicted to be involved in regulation of transcription by RNA polymerase II and retinal rod cell development and to act upstream of or within positive regulation of gene expression and positive regulation of transcription by RNA polymerase II. Predicted to be located in cytosol and nucleoplasm and to be active in nucleus. |
| ***Gngt1*** | G protein subunit gamma transducin 1 | NM_001135777 | Predicted to enable G-protein beta-subunit binding activity. Acts upstream of or within cardiac muscle cell apoptotic process and cellular response to hypoxia. Located in photoreceptor inner segment and photoreceptor outer segment. Part of heterotrimeric G-protein complex. |
|  | **<u>RPE MARKERS</u>** |  |  |
| ***Rpe65*** | retinoid isomerohydrolase RPE65 | NM_053562 | Enables all-trans-retinyl-ester hydrolase, 11-cis retinol forming activity. Involved in circadian rhythm; neural retina development; and retinoid metabolic process. Located in cell body. |
| ***Tjp1*** | tight junction protein 1 | NM_001106266 | Enables protein C-terminus binding activity. Involved in cellular response to glucose stimulus; negative regulation of vascular permeability; and response to lipopolysaccharide. Located in bicellular tight junction; cytoplasm; and gap junction. |
| ***Best1*** | bestrophin 1 | NM_001011940 | Predicted to enable identical protein binding activity; to contribute to chloride channel activity; to be involved in transepithelial chloride transport; to act upstream of or within chloride transport; detection of light stimulus involved in visual perception; and regulation of calcium ion transport. Predicted to be located in basolateral plasma membrane. Predicted to be part of chloride channel complex. |
|  | **<u>HOUSEKEEPER</u>** |  |  |
| ***Hmbs*** | hydroxymethylbilane synthase | NM_013168 | Part of the hydroxymethylbilane synthase superfamily in which the gene product is the third enzyme in the heme biosynthetic pathway and serves as a catalyst for the head to tail condensation of four molecules of porphobilinogen into the linear hydroxymethylbilane. |

#### Statistical Analysis

Statistical analysis was performed using Graphpad Prism, using t-tests (paired and unpaired), Mann-Whitney U, one-way ANOVA for multiple comparisons and/or Tukey post-hoc analysis. The significance level was set at p<0.05.

## Results

### Intraperitoneal HNG Injection Studies

Animals were weighed prior to each injection (HNG or saline) and any side effects were recorded. No side effects were observed throughout the study. No change in behavior was found. We found no significant changes in the weights between the saline and HNG (low or high dose) groups at the end of the study (**Figure 1**).

**Figure 1.**
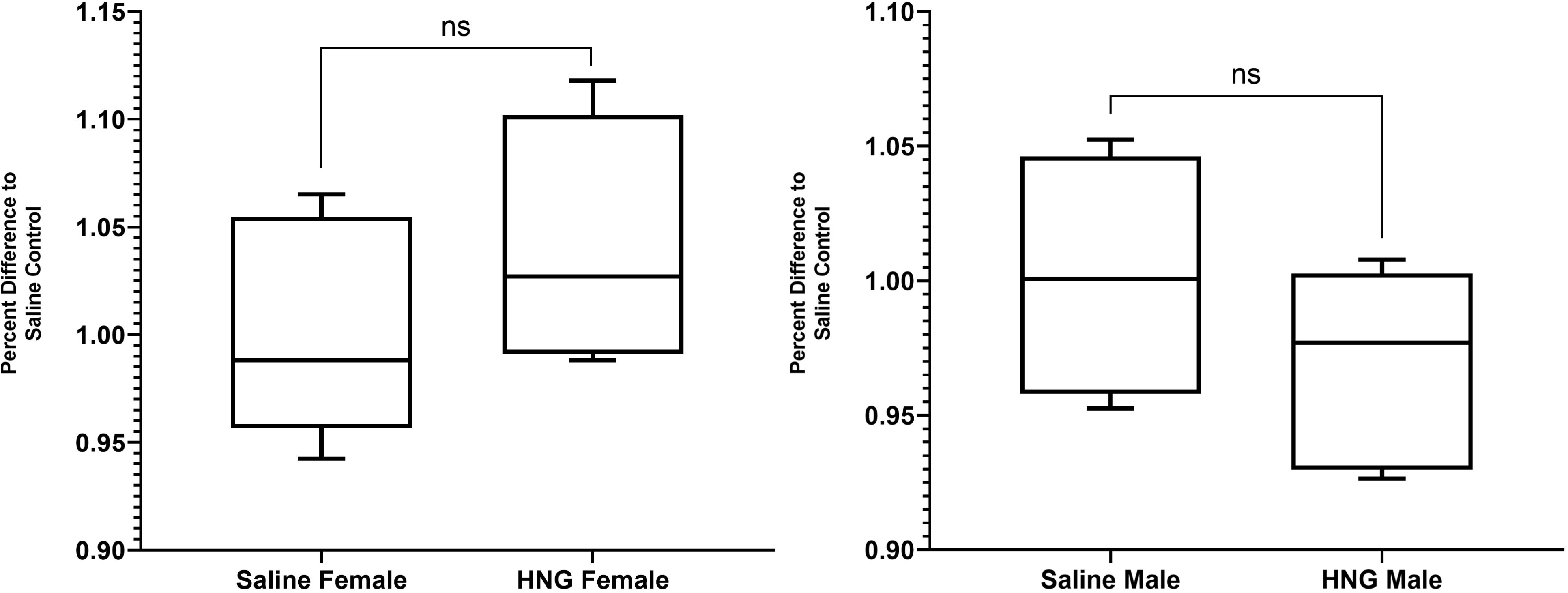
No change in animal weight at 4 WAFI.

### Retinal Gene Expression Levels after Intraperitoneal HNG Injection

#### Gene Expression for IP Low Dose HNG (0.4mg/kg) at 1 WAFI

The RPE showed a significant increase in expression of *Tnf*α (1.789-fold P=0.0023) and a significant decrease in expression of *Rlbp1* (0.6781-fold, P=0.0271) in animals treated with low dose HNG for one week (**Supplemental Table 1**). There were no statistically significant changes in gene expression in the neuroretina of animals treated with low dose HNG for one week.

#### Gene Expression for IP Low Dose HNG (0.4mg/kg) at 4 WAFI

The neuro-retina showed decreased levels of proapoptotic *Ddit3* (0.7658-fold, P=0.0159) and increased levels of Crx (1.468-fold, P=0.0357) and *Tjp1* (1.567-fold, P=0.0286) (**Figure 2**, **Supplemental Table 2** The RPE cells demonstrated no significant changes in gene expression.

**Figure 2.**
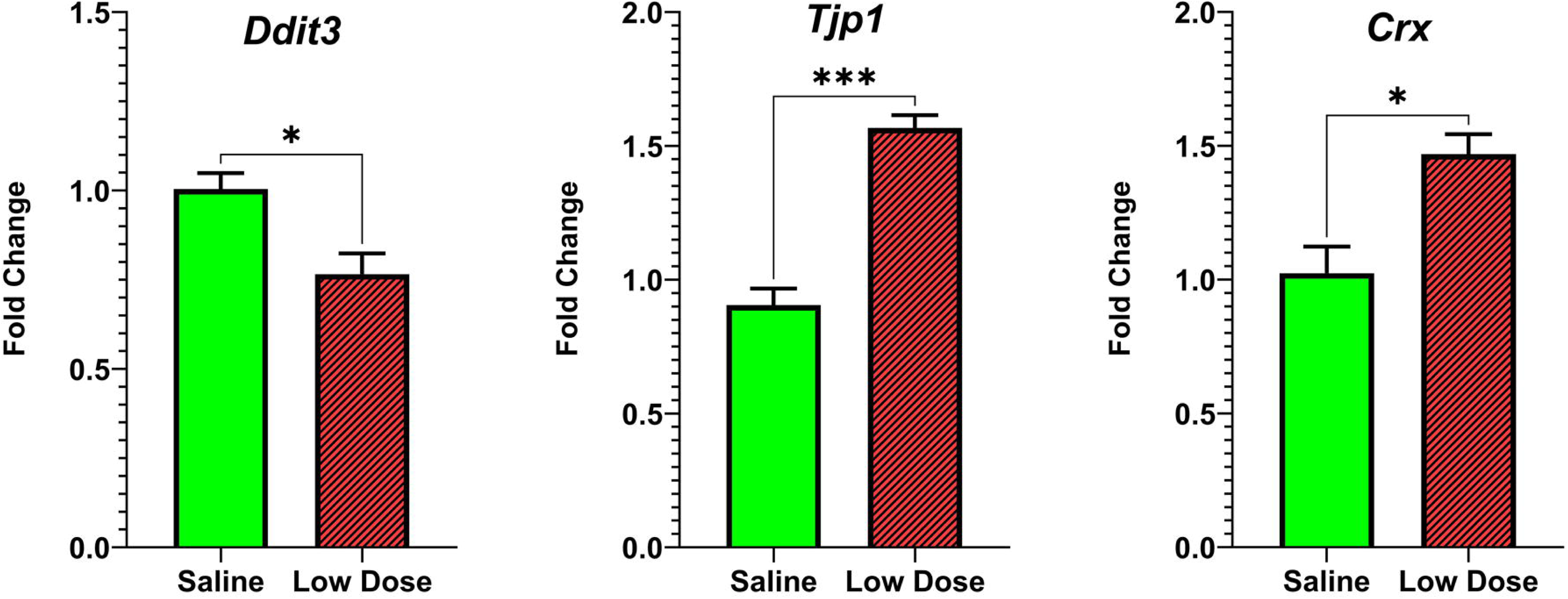
Effect of Low dose (0.4mg/kg) HNG on gene expression in neuroretina 4 WAFI. Expression of *Ddit3* was significantly decreased and expression of *Tjp1* and *Crx* were significantly increased 4 WAFI with low dose HNG compared with saline in the neuroretina

#### Gene Expression for IP High Dose HNG (4mg/kg) at 1 WAFI

In the RCS rats treated with High Dose HNG (n=9), the RPE showed decreased expression levels of apoptotic genes (Bcl2l1, 0.587-fold, P=0.0190) and *Casp7* (0.5737-fold, P=0.0321) compared to the saline treated samples (n=9) (**Figure 3A**, **Supplemental Table 3**) The neuro-retina showed increased levels of Crx (1.309-fold, P= 0.0103) (**Figure 3B)**. There were no statistically significant changes in other genes’ expression.

**Figure 3.**
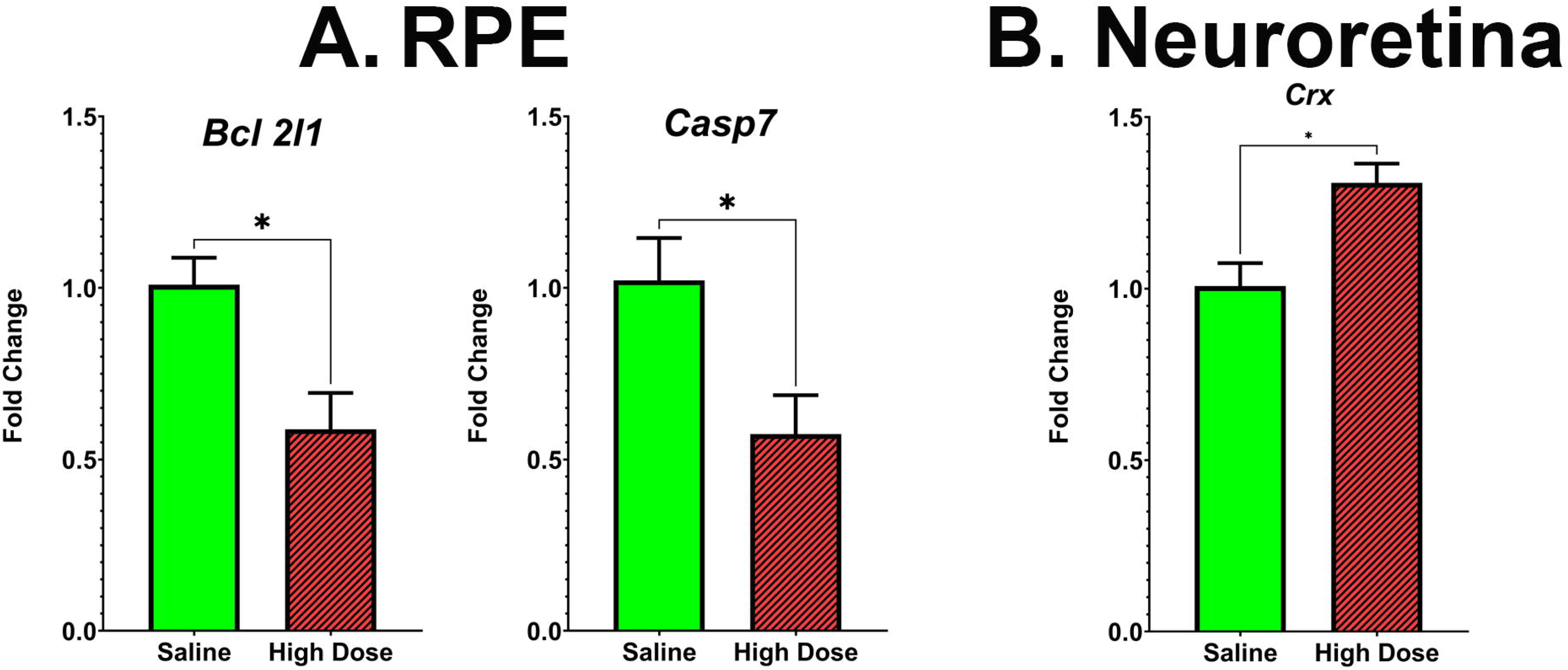
Effect of High dose (4.0mg/kg) HNG on gene expression in the RPE and Neuroretina 1 WAFI. (a) Expression of Bcl2l1 and Casp7 were both significantly decreased 1 WAFI with high dose HNG compared with saline in the RPE. (b) Expression of Crx was significantly increased 1 WAFI with high dose HNG compared with saline in the Neuroretina.

#### Gene Expression for IP High Dose HNG (4mg/kg) at 4 WAFI

The RPE showed increased expression levels of pro-apoptotic genes (*Ddit3*, 1.842-fold, P=0.0317; *Casp3*,1.65-fold, P=0.0079); inflammation genes (*Il6*, 1.782-fold, P=0.0286); antioxidant genes (Sod2, 1.797-fold, 0.0286); and RPE genes important for normal function (*Best1*, 1.698-fold, P=0.0317 (**Figure 4a**, **Supplemental Table 4**). There was also a lower expression level of *E2f1* (0.569-fold, P=0.016) in the RPE. The neuroretina showed increased levels of proapoptotic *Casp7* (1.309-fold, P=0.0159) (**Figure 4b**, **Table 4**). There were no statistically significant changes in the expression of other genes.

**Figure 4.**
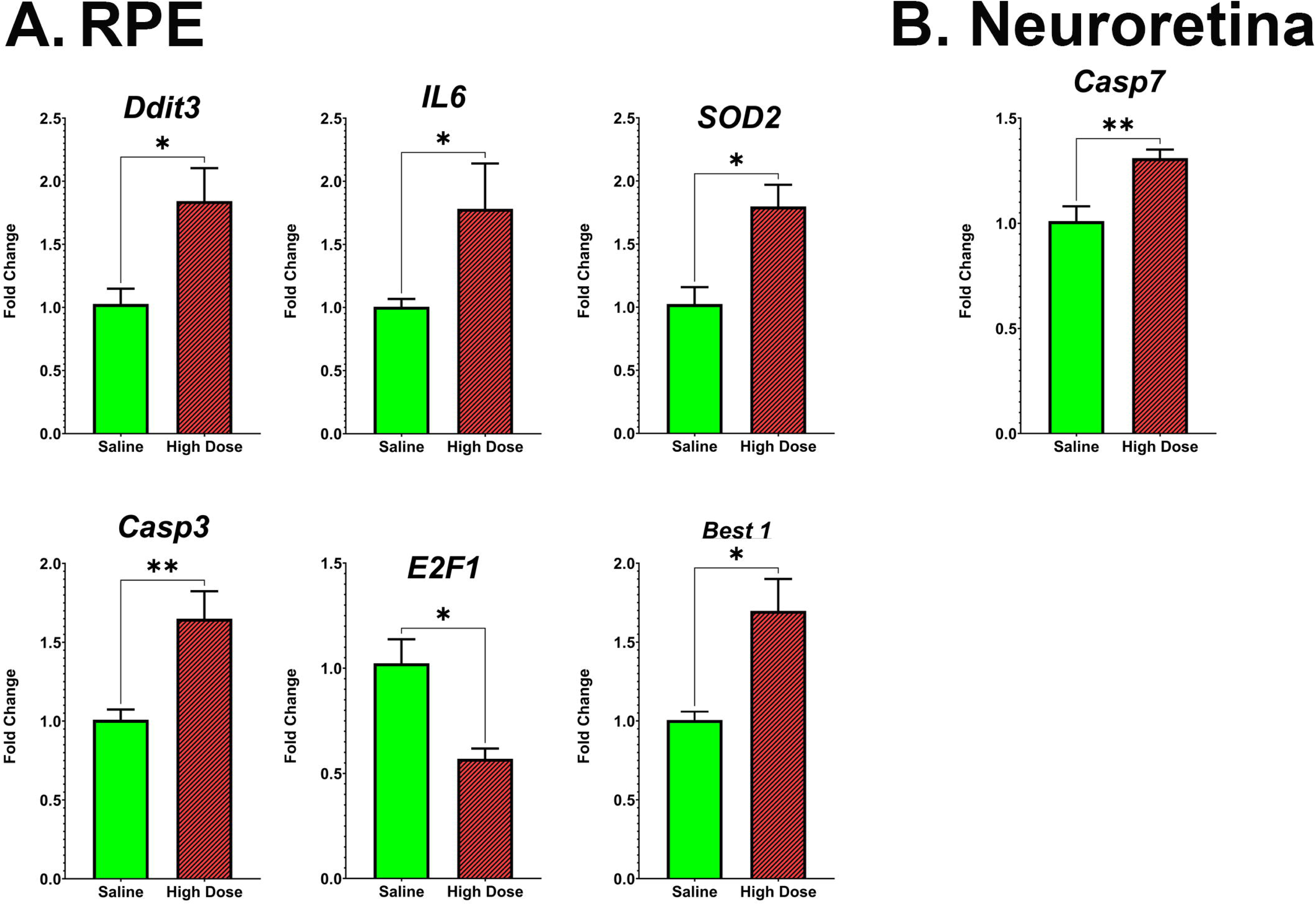
Effect of High dose (4.0mg/kg) HNG on gene expression in the RPE and Neuroretina 4 WAFI. (a) Expression of *Ddit3*, *Il6*, *Sod2*, *Casp3*, and *Best1* were all significantly increased, and *E2f1* were significantly decreased 4 WAFI with high dose HNG in the RPE. (b) Expression of *Casp7* was significantly increased 4 WAFI with high dose HNG in the neuroretina.

#### Results of ERG at 1 and 4 WAFI

The ERG analyses were performed first on Low Dose HNG (0.4mg/kg), and Sham-Saline at 4 WAFI. There was no difference between Low Dose HNG and Saline in a-wave and b-wave of scotopic and photopic full field ERG (**Figure 5**). Later, High Dose HNG (4mg/kg) and Saline groups were tested with ERG. At 1 and 4 WAFI, there was no significant difference in scotopic and photopic a-wave (data not shown), or scotopic and photopic b-wave between High dose HNG and Sham-Saline groups (**Figure 5**).

**Figure 5.**
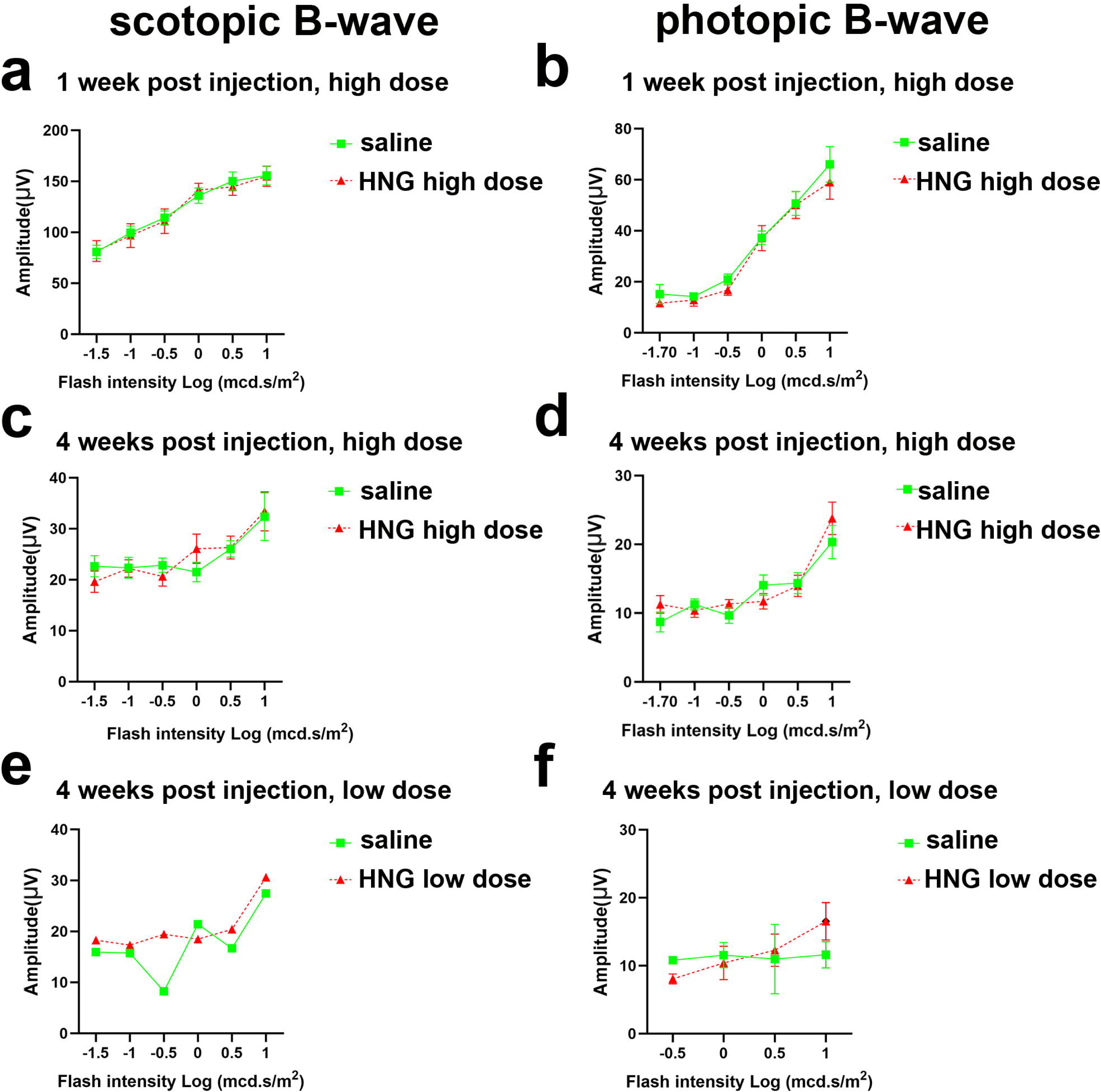
ERG Results from low (0.4mg/kg) and high dose (4mg/kg) IP Injections at 1 and 4 WAFI. No significant change was found.

#### Visual function improvement evaluated by optokinetic testing (OKT)

At 4 WAFI, optokinetic testing in RCS rats showed that the visual acuity of eyes with high dose HNG (n=11) showed significant improvement from that of saline groups (n=9. P<0.05). (**Figure 6**) There was no significant difference in any other time points or doses tested.

**Figure 6.**
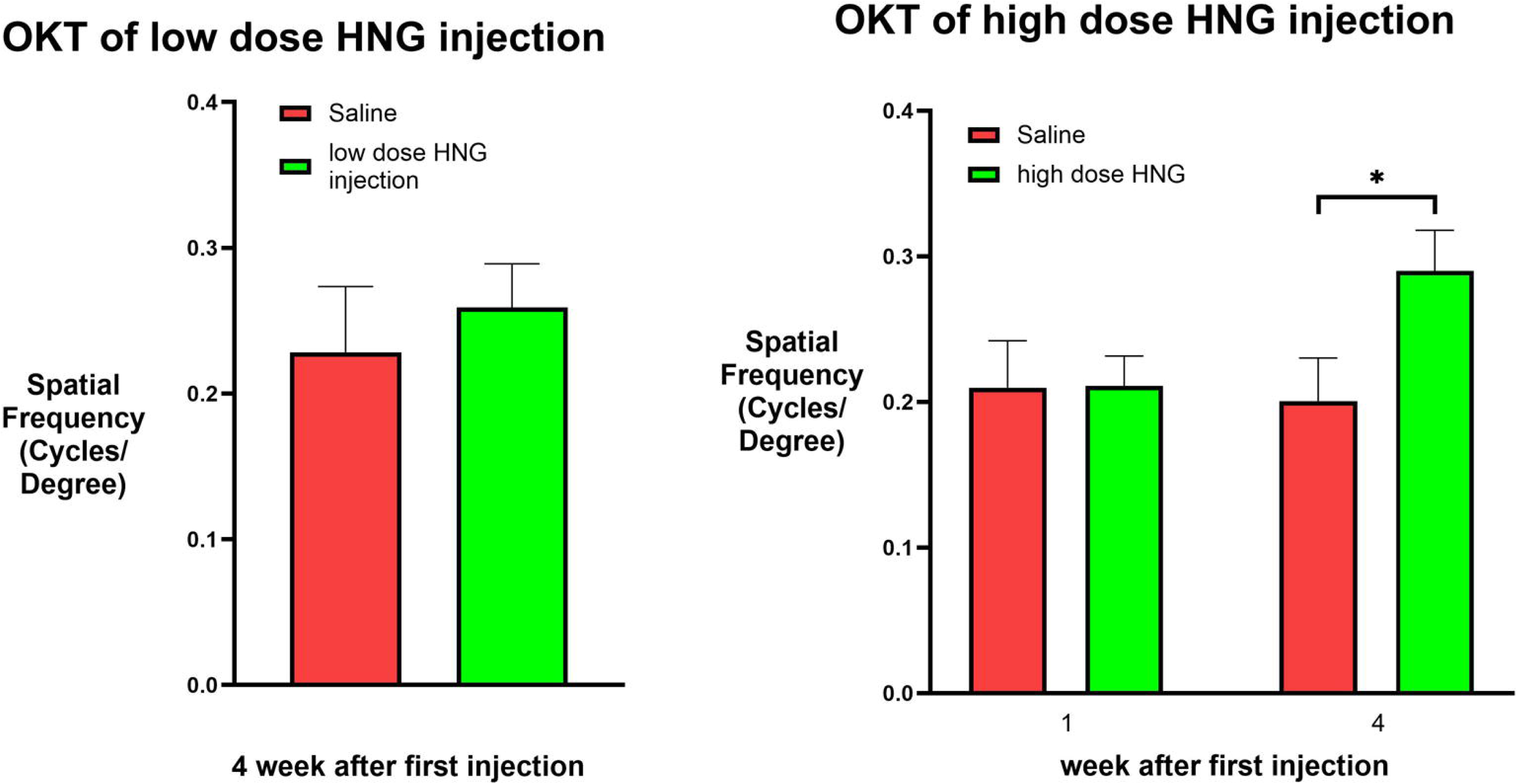
Effect of high dose IP injections of HNG on OKT at both 1 WAFI and 4 WAFI. OKT was improved at 4 WAFI. Asterisk indicates significant difference at P>0.05.

## Discussion

HNG has demonstrated cytoprotective effects in human cybrid cell lines and animal models of neurodegeneration. In a study evaluating an analog of Humanin in ameliorating streptozotocin-induced diabetic nephropathy in Sprague Dawley rats, [S14G]-humanin was administered intra-peritoneal (once daily for a course of sixteen weeks) at a dosage of 0.4 mg/Kg of body weight ^45^. Another study reported that humanin exhibits neuroprotective effects in vitro in human cell culture models and enhances cognition in vivo in aged mice (Kelvin Yen 2018). In the present study, we used intraperitoneal injections to deliver either low dose (0.4 mg/kg) or high dose (4mg/kg) HNG to the RCS rats and showed that HNG IP injections can modulate in RPE and neuroretina gene expression levels and in the short-term, improve retina function in the RCS rat model.

### Low Dose HNG

#### RPE

The Low dose IP injections at 1 WAFI resulted in upregulation at *Tnf*α and downregulation of Rlbp1 in the RPE cells, whereas by 4 WAFI, no significant changes in gene expression were observed across all pathways. In a rat model of ulcerative colitis, intraperitoneal HNG (10 or 20 µM) reduced expression of *Tnf*α and *Il-1*β and decreased caspase 3 activities^46^. Similarly, in a rat model of pituitary tumors, 5 µM HN inhibited the proapoptotic effect of *Tnf*α on cultured anterior pituitary cells^47^. In ApoE deficient mice on a high-cholesterol diet, after 16 weeks of intraperitoneal HN (4 mg/kg/day) treatment, *Tnf*α, MCP-1 and osteopontin were downregulated, along with decreased apoptosis, compared to the untreated ApoE-/- mice ^48^. Similar decline in the pro-inflammatory *Tnf*α and *IL-1*β were seen after HN treatment in HUVEC culture ^49^.

TNF-α is a key pivotal mediator of inflammation in AMD, known for disrupting endothelial integrity and initiating inflammatory pathways that accelerate disease progression ^50,51^. However, studies assessing TNF-α level in aqueous humour or in serum remain inconclusive ^52–55^. Although TNF-α is theoretically expected to be elevated in AMD, most comparisons between AMD and control groups have shown no significant differences ^53,55^. furthermore, TNF-α was significantly higher in patients who improved than in those who deteriorated ^53^.

Overall, systemic and local measurements of TNF-α remain inconsistent. While TNF-α inhibition is effective for some ocular diseases such as uveitis ^56^, clinical outcomes for anti–TNF-α therapy in AMD, whether systemic ^57^ or intravitreal ^58,59^, have been variable. Future studies should clarify these discrepancies.

Our observation of elevated TNFα at 1 WAFI may reflect either the uncertain role of TNFα or the short duration and low dose of treatment, particularly since no change was observed at 4 WAFI. Further investigation is warranted.

RLBP1 encodes cellular retinal-binding protein (CRALBP), an 11-cis-retinal–binding protein expressed in Müller and RPE cells ^60^, and is essential for the visual cycle. RLBP1 mutations cause three clinical subtypes—Bothnia dystrophy, retinitis punctata albescens, and Newfoundland rod-cone dystrophy ^61^. Gene therapy using AAV vectors expressing RLBP1 has improved the visual cycle in Rlbp1−/− mice ^62^ and in patients with RLBP1-associated retinal dystrophy ^63^. We observed downregulation of Rlbp1 in RPE cells at 1 WAFI but not at 4 WAFI. Further studies are needed.

#### Neuro-retina

Interestingly, at 1 WAFI, no gene expression changes were seen for any of the neuroretinal genes, suggesting that either higher HNG doses or a longer exposure is required to modulate the neuroretinal genes. By 4 WAFI, *Ddit3 was significantly downregulated* (p=0.0159), while *Tjp1* (p=0.0286) and *Crx* (p=0.0357) were upregulated. *The DNA damage inducible transcript 3 (DDIT3, also known as transcription factor C/EBP-homologous protein, CHOP) is an ER stress effector that promotes* ER stress and the subsequent inflammation and apoptosis induced by lipopolysaccharide (LPS) exposure ^64,65^.

#### DDIT3 is implicated in neurodegeneration

Human RPE cybrids harboring AMD mitochondria show markedly elevated *DDIT3* (also known as CHOP, 633.9%), *Caspase-3* (125.7%). *Caspase-7* (181.3%) levels, with reduced *E2F1* (66.2%) and *SOD2* levels (23.1%) compared to the cybrids with age-matched normal mitochondria. HNG treatment significantly reduces these pro-apoptotic and ER stress markers^25^. In primary human RPE cell cultures, HN pretreatment decreases ER-stress induced apoptosis by elevating mitochondria glutathione and lowering CHOP (DDIT3) levels ^30^. Similarly, Sreekumar et al. demonstrated that HN localizes to RPE cells and protected against oxidative stress by improving mitochondrial biogenesis and bioenergetics^16^. The observed reduction of Ddit3 expression by HNG in our study further supports DDIT3’s role in retinal degeneration and highlights HNG’s therapeutic potential.

Tight junction protein-1 (TJP1), also known as the tight junction marker ZO-1, is a peripheral membrane phosphoprotein of the zonula occludens family ^66^ and a key component linking junctional proteins to the cytoskeleton to maintain epithelial integrity ^67^. Reduced ZO-1 expression in RPE cells is associated with epithelial–mesenchymal transition and compromised retinal support ^67,68^, leading to pathologies such as age-related macular degeneration (AMD) and diabetic retinopathy ^67,68^. Thus, our data, HNG-induced upregulation of Tjp1, *suggests its potential to preserve RPE integrity and treat AMD*.

Crx (cone–rod homeobox) is an transcription factor critical for photoreceptor development and differentiation, regulating numerous phototransduction and metabolic genes ^69^. Dysregulated Crx–Otx2 interaction contributes to early-onset retinal degeneration ^70^. Mutations in upstream regulators of Crx–Otx2 disrupt gene activation balance, causing aberrant photoreceptor gene expression and apoptosis ^70^. Crx also partners with Nrl and Nr2e3 to promote rod-specific gene expression while suppressing cone genes during differentiation ^69^. Loss or mutation of Crx results in photoreceptor degeneration and disorders such as Leber congenital amaurosis (LCA) and cone–rod dystrophy ^71^. In our study, HNG-induced upregulation of Crx further supports its potential as a therapeutic agent for AMD.

### High Dose HNG

As expected, high-dose HNG altered the expression of more genes than low-dose HNG.

#### RPE

At 1 WAFI, HNG-treated RPE cells showed downregulation of the pro-apoptotic gene Casp7 (p = 0.0321) and the anti-apoptotic Bcl2l1 (p = 0.0189). While the latter appears inconsistent with HNG’s beneficial effects, Bcl2l1 has multiple cellular roles. BCL2L1 encodes the anti-apoptotic protein BCL-XL ^72^, which provides a strong selective advantage to hPSCs under stress conditions such as thawing, expansion, and cloning^72^. BCL-XL localizes mitochondria to prevent cytochrome C release and caspase-dependent apoptosis through interactions with other BCL-2 family members ^73,74^. Beyond its anti-apoptotic role, BCL-XL contributes to metabolism, mitochondrial dynamics, and calcium homeostasis ^73,74^, and has been implicated in regulating cell fate by inhibiting neuroectodermal differentiation ^75^. In RPE tissue and ARPE19 cells, BCL-XL supports cell survival ^76^, but persistent expression in senescent RPE cells contributes to apoptosis resistance and tissue dysfunction in aging and AMD models ^75,77,78^. Inhibiting BCL-XL in senescent RPE is therefore a promising therapeutic approach for AMD ^77,78^. Given its roles in mitochondrial and calcium homeostasis—critical to RPE physiology—the downregulation of Bcl2l1 by HNG may also contribute to its protective effects.

At 4 WAFI, high-dose HNG altered more genes than the low dose (6 vs 0) in RPE cells, including upregulation of pro-apoptotic (Casp3), ER stress (Ddit3), inflammatory (Il-6), antioxidant (Sod2), and RPE marker (Best1) genes, while E2f1 expression was significantly reduced. E2F1 is a transcription factor regulating genes involved in DNA replication, repair, the cell cycle, and apoptosis ^79^. Its biological functions are modulated by post-translational modifications ^80^. IL-6 is a pleiotropic cytokine that primarily acts as a pro-inflammatory factor ^81^. In RPE cells, IL-6 inhibits Sirt1 through PI3K/AKT/mTOR-mediated phosphorylation, suppressing the E2F1/HMGA1/G6PD pathway and thereby increasing oxidative stress and cell death ^82^. Although HNG increased Casp3, Ddit3, and Il-6 while reducing E2f1, these changes may reflect alternative functions and warrant further investigation.

Superoxide dismutases (SODs) are critical antioxidant enzymes that prevent oxidative stress by metabolizing reactive oxygen species (ROS) ^83^. SOD2, localized in mitochondria, detoxifies superoxide radicals generated during oxidative phosphorylation at complexes I and III, converting them into hydrogen peroxide and oxygen ^84^. Upregulation of Sod2 reduces oxidative stress, supporting mitochondrial protection, a key factor in neurodegenerative disease mitigation ^85^. Thus, HNG-induced Sod2 upregulation suggests a potential therapeutic mechanism.

The Best gene family encodes Ca²⁺-activated anion channels with diverse physiological roles in multiple organs, including the eye ^86^. Best1 is predominantly expressed in RPE cells and is genetically associated with various retinal degenerations ^87,88^, encompassing over 350 known mutations leading to progressive vision loss and blindness ^89,90^. Given Best1’s critical function in retinal health and its link to untreatable vision disorders, the observed Best1 upregulation by HNG further supports its therapeutic potential in retinal degenerative diseases.

#### Neuro-retina

At 1 WAFI, the neuroretina exhibited elevated *Crx* expression (p=0.0103), while other genes remained comparable to saline-treated controls. As described above, *Crx* regulates multiple photoreceptor-specific genes essential for photoreceptor development and differentiation. Its upregulation supports the beneficial effect of HNG in retinal degeneration (RD) diseases. At 4 WAFI, *Casp7* expression increased (p=0.0159), whereas other genes remained unchanged relative to controls. Overall, more genes were modulated in RPE cells (six genes) than in neuroretina (one gene), likely due to the reduced neuroretinal thickness in RCS rats by 30 days or a greater sensitivity of RPE cells to HN-mediated protection. Sreekumar et al. reported that polarized human RPE cells contain high intracellular HN co-localized with mitochondria ^16^. Exogenous FITC-labeled HN peptides showed robust uptake and mitochondrial translocation in RPE cells compared to controls, indicating a mitochondria-targeted protective role. Moreover, RPE cells express all three HN receptors—ciliary neurotrophic factor receptor (CNTFRα), transmembrane glycoprotein gp130 (GP130), and cytokine receptor WSX1 ^16,30^—which may account for the stronger HNG effects observed in RPE cells than in neuroretina.

#### Functional Testing with High Dose HNG

No significant differences were observed in scotopic or photopic a- and b-wave amplitudes between high-dose, low-dose HNG, and sham-saline groups. The absence of detectable ERG changes at 1 WAFI may reflect the single early injection and the time required for HNG to exert measurable effects. By 4 WAFI, the rapid retinal degeneration characteristic of the RCS rat resulted in undetectable ERG responses across all groups; therefore, a potential beneficial effect of HNG cannot be excluded, as ERG may lack the sensitivity to detect subtle improvements. Interestingly, while no improvement in OKT was seen at 1 WAFI following high-dose intraperitoneal HNG, visual function improved at 4 WAFI. Molecular analyses revealed increased expression of several genes in both neuroretina and RPE tissues, suggesting that intraperitoneally administered HNG may cross the blood–retina barrier or induce systemic changes affecting these tissues.

Overall, intraperitoneal HNG administration in the RCS rat model (a) modulated gene expression in RPE and neuroretina, indicating possible blood–retina barrier penetration or systemic regulation, and (b) improved visual performance at 4 WAFI as measured by OKT. Given the variability in molecular responses and the limited clinical applicability of intraperitoneal delivery for retinal diseases, future studies will focus on evaluating alternative HNG delivery methods in this model of retinal degeneration.

## Acknowledgments

The authors want to acknowledge the contribution of Dr. M.C. Kenney, UCI (who passed away in December 2023) to the initiation, funding and completion of this project. The authors would like to acknowledge the technical assistance of Shari R. Atilano, Mithalesh Singh (Kenney lab), and Seiler lab staff members Robert Sims, Karla Echeverria, Jay Santoso, Reeva Reyes, Jeffrey Delgado; and students Alice Avetyan, Candice Wu, Caroline Lee, Jennie Xu, Kevin Wu, Angel Blanquel, Adeline Cheng, Winnie Luong, Julianne Frances Agapinan, Ceci Zhang, Ani Petrosyan, Ani Khachigian, Ryan Pavey. Christopher Quimpo, Maihan Phan, and other students. This research was funded by the National Eye Institute, grant R01 EY027363 and to a small part with grant R01 EY031834. The authors acknowledge support to the Gavin Herbert Eye Institute at the University of California, Irvine from an unrestricted grant from Research to Prevent Blindness and from NIH core grant P30 EY034070.

## Author contributions

BL: performed experiments, collected & analyzed data; wrote the paper; approved the final version

KS: performed experiments, collected & analyzed data; wrote the paper; approved the final version

MO: performed experiments, collected & analyzed data; wrote the paper; approved the final version

NI: performed experiments, collected data; approved the final version

MJS: performed experiments, analyzed data; wrote the paper; edited figures; provided funding; approved the final version

---

**Supplemental Table 1.** qPCR Results from low dose (0.4mg/kg) IP Injections 1 WAFI (RPE and Neuroretina)

**Supplemental Table 2.** qPCR Results from low dose (0.4mg/kg) IP Injections 4 WAFI (RPE and Neuroretina)

**Supplemental Table 3.** qPCR Results from high dose (4mg/kg) IP Injections 1 WAFI (RPE and Neuroretina)

**Supplemental Table 4**. qPCR Results from high dose (4mg/kg) IP Injections 4 WAFI (RPE and Neuroretina)

