## Supplemental tables for "Neuroprotective Effect of Intraperitoneal Humanin-G in Retinal Degeneration of Royal College of Surgeons Rats"

**Supplemental Table 1**: qPCR Results from low dose (0.4mg/kg) IP Injections 1 WAFI (RPE and Neuroretina)

|  | **RPE** |  |  |  | **Retina** |  |
| --- | --- | --- | --- | --- | --- | --- |
| **Gene** | **HNG Treated Fold Expression (compared to saline)** | **p-value** |  | **Gene** | **HNG Treated Fold Expression (compared to saline)** | **p-value** |
| *Bax* | 1.091 | 0.6224 |  | *Bax* | 0.9213 | 0.5422 |
| *Bcl2l13* | 1.138 | 0.3002 |  | *Bcl2l13* | 1.024 | 0.8706 |
| *Casp3* | 1.241 | 0.2565 |  | *Casp3* | 1.131 | 0.5471 |
| *Ddit3* | 1.23 | 0.5288 |  | *Ddit3* | 1.085 | 0.5127 |
| *Il1b* | 0.8221 | 0.4262 |  | *Il1b* | 0.8178 | 0.3683 |
| *Tnfα* | **1.789** | **0.0023**** |  | *Tnfα* | 1.028 | 0.7416 |
| *Sod2* | 1.046 | 0.7931 |  | *Sod2* | 0.9722 | 0.7528 |
| *Best1* | 1.268 | 0.4062 |  | | | |
| *Rpe65* | 0.8794 | 0.5632 |  |  |  |  |
| *Rlbp1* | **0.6781** | **0.0271*** |  |  |  |  |

**Supplemental Table 2**: qPCR Results from low dose (0.4mg/kg) IP Injections 4 WAFI (RPE and Neuroretina)

|  | **RPE** |  |  |  | **Retina** |  |
| --- | --- | --- | --- | --- | --- | --- |
| **Genes** | **Low Dose IP HNG Fold expression compared with saline** | **p-value** |  | **Genes** | **Low Dose IP HNG Fold expression compared with saline** | **p-value** |
| *Ddit3* | 1.283 | 0.2286 |  | *Ddit3* | **0.7658** | **0.0159 (*)** |
| *Il6* | 0.66 | 0.4 |  | *Il6* | 0.6841 | 0.25 |
| *Il1b* | 1.306 | 0.8571 |  | *Il1b* | 1.196 | 0.2857 |
| *Sod2* | 1.413 | 0.2286 |  | *Sod2* | 1.035 | >0.9999 |
| *Tnfα* | 0.8782 | 0.6286 |  | *Tnfα* | 1.49 | 0.25 |
| *Bax* | 1.03 | 0.8571 |  | *Bax* | 0.8223 | 0.2857 |
| *Bcl2l1* | 0.8567 | 0.4 |  | *Bcl2l1* | 1.031 | >0.9999 |
| *Bcl2l13* | 1.077 | 0.8571 |  | *Bcl2l13* | 1.169 | 0.5556 |
| *Casp3* | 0.9553 | 0.8571 |  | *Casp3* | 1.018 | 0.9048 |
| *Casp7* | 0.8658 | 0.1143 |  | *Casp7* | 0.9767 | >0.9999 |
| *Casp9* | 0.8867 | 0.6286 |  | *Casp9* | 1.049 | 0.9048 |
| *Ccl2* | 1.646 | 0.8571 |  | *Ccl2* | 1.317 | 0.4127 |
| *E2f1* | 1.211 | 0.8571 |  | *E2f1* | 1.238 | 0.0635 |
| *Tjp1* | 1.308 | 0.1143 |  | *Tjp1* | **1.567** | **0.0286 (*)** |
| *Best1* | 1.707 | 0.2286 |  | *Crx* | **1.468** | **0.0357 (*)** |
| *Rpe65* | 1.044 | >0.9999 |  | *Gngt1* | 1.089 | 0.9048 |
|  |  |  |  | *Nrl* | 1.653 | 0.9048 |
|  | | | | *Rom1* | 0.7292 | 0.4127 |
|  |  |  |  | *Hspa5* | 1.011 | 0.9048 |

**Supplemental Table 3**: qPCR Results from high dose (4mg/kg) IP

Injections 1 WAFI (RPE and Neuroretina)

|  | **RPE** |  |  |  | **Neuroretina** |  |
| --- | --- | --- | --- | --- | --- | --- |
| **Genes** | **High Dose (4mg/kg) HNG IP Fold expression compared with saline** | **p-value** |  | **Genes** | **High Dose (4mg/kg) HNG IP Fold expression compared with saline** | **p-value** |
| *Ddit3* | 1.209 | 0.2119 |  | *Ddit3* | 1.074 | 0.6873 |
| *Il1b* | 1.302 | 0.3226 |  | *Il6* | 1.137 | 0.6369 |
| *Sod2* | 0.966 | 0.8133 |  | *Il1b* | 1.448 | 0.1967 |
| *Tnfα* | 1.19 | 0.8023 |  | *Sod2* | 1.038 | 0.8013 |
| *Bax* | 1.116 | 0.8431 |  | *Tnfα* | 1.493 | 0.1407 |
| *Bcl2l1* | **0.5877** | **0.0189 (*)** |  | *Bax* | 1.071 | 0.7491 |
| *Bcl2l13* | 0.7626 | 0.3433 |  | *Bcl2l1* | 1.313 | 0.3101 |
| *Casp3* | 0.6843 | 0.1908 |  | *Bcl2l13* | 1.002 | 0.985 |
| *Casp7* | **0.5737** | **0.0321 (*)** |  | *Casp3* | 1.148 | 0.0835 |
| *Casp9* | 0.8067 | 0.4281 |  | *Casp7* | 1.085 | 0.241 |
| *Ccl2* | 0.789 | 0.3173 |  | *Casp9* | 1.059 | 0.4636 |
| *E2f1* | 1.063 | 0.8834 |  | *Ccl2* | 2.451 | 0.2405 |
| *Tjp1* | 0.9617 | 0.7282 |  | *E2f1* | 1.023 | 0.8115 |
| *Best1* | 1.096 | 0.9697 |  | *Tjp1* | 1.132 | 0.3359 |
| *Rpe65* | 1.054 | 0.8924 |  | *Crx* | **1.309** | **0.0103 (*)** |
| *Hspa5* | 0.9845 | 0.8186 |  | *Gngt1* | 0.9837 | 0.7516 |
|  |  |  |  | *Nrl* | 1.271 | 0.3044 |
|  | | | | *Rom1* | 1.239 | 0.6064 |
|  |  |  |  | *Hspa5* | 1.334 | 0.3237 |

**Supplemental Table 4**: qPCR Results from high dose (4mg/kg) IP Injections 4 WAFI(RPE and Neuroretina)

|  | **RPE** |  |  |  | **Retina** |  |
| --- | --- | --- | --- | --- | --- | --- |
| **Genes** | **High Dose (4mg/kg) HNG IP Fold expression compared with saline** | **p-value** |  | **Genes** | **High Dose (4mg/kg) HNG IP Fold expression compared with saline** | **p-value** |
| *Ddit3* | **1.842** | **0.0317 (*)** |  | *Ddit3* | 1.03 | >0.9999 |
| *Il6* | **1.782** | **0.0286 (*)** |  | *Il6* | 1.611 | 0.3095 |
| *Il1b* | 1.028 | 0.9048 |  | *Il1b* | 1.399 | 0.0635 |
| *Sod2* | **1.797** | **0.0286 (*)** |  | *Sod2* | 1.102 | 0.6905 |
| *Tnfα* | 1.137 | 0.7302 |  | *Tnfα* | 1.307 | 0.0952 |
| *Bax* | 1.233 | 0.1111 |  | *Bax* | 1.04 | 0.8413 |
| *Bcl2l1* | 0.8425 | 0.4127 |  | *Bcl2l1* | 0.9644 | 0.5476 |
| *Bcl2l13* | 0.8026 | 0.25 |  | *Bcl2l13* | 1.033 | >0.9999 |
| *Casp3* | **1.65** | **0.0079 (**)** |  | *Casp3* | 1.149 | 0.4206 |
| *Casp7* | 1.062 | 0.6905 |  | *Casp7* | **1.309** | **0.0159 (*)** |
| *Casp9* | 1.059 | 0.6905 |  | *Casp9* | 1.337 | 0.0556 |
| *Ccl2* | 1.977 | 0.2222 |  | *Ccl2* | 1.156 | 0.4206 |
| *E2f1* | **0.5693** | **0.0159 (*)** |  | *E2f1* | 1.023 | >0.9999 |
| *Tjp1* | 0.84 | 0.4206 |  | *Tjp1* | 0.9477 | 0.3095 |
| *Best1* | **1.698** | **0.0317 (*)** |  | *Crx* | 0.8771 | 0.3095 |
| *Rpe65* | 1.083 | >0.9999 |  | *Gngt1* | 0.9883 | 0.6905 |
| *Hspa5* | 0.8088 | 0.1508 |  | *Nrl* | 0.9232 | 0.6905 |
|  | | | | *Rom1* | 0.9936 | >0.9999 |
|  |  |  |  | *Hspa5* | 0.9206 | 0.3095 |
